# Determining the Role of Environmental Covariates on Planktivorous Elasmobranch Population Trends within an Isolated Marine Protected Area

**DOI:** 10.1101/2022.09.28.509935

**Authors:** Julia Saltzman, Easton R. White

## Abstract

Several studies have found predictable relationships between the behavior of planktivores and environmental conditions, suggesting that planktivores may be especially sensitive to environmental change. However, many studies to date are based on limited observations, include few of the many environmental covariates which could influence planktivores, and do not occur over long enough time periods to make inferences about the potential effects of environmental change. As such, long term datasets on planktivores are necessary to disentangle the potential impacts of oceanographic and environmental variability. In this study, to elucidate the relationship between plankivores and environmental variability, we use data obtained over the last 28 years by a small group of divemasters at Cocos Island, Costa Rica, one of the oldest marine reserves in the world. We found that, in general, for planktivorous elasmobranchs, several environmental variables, such as, chlorophyll A, lunar cycle, and salinity have clear influences on their occurrence and relative abundances. We found that in the phases of lower illuminations, there were significant increases in abundance of mobula rays. Specifically, a 0.10 *mg*/*m*^3^increase in Chlorophyll A correlated with 26% decrease in whale sharks. We found that increases in salinity correlated with increases in mobula abundance but did not correlate with observations of mantas or whale sharks. We also found that omission of environmental covariates can lead to overprediction and underprediction of relative abundances. Our findings highlight the need to take environmental conditions into account when evaluating the efficacy of marine protection.

## 1. Introduction

Planktivorous elasmobranchs are highly influenced by environmental variability and food availability. Unlike species and groups of elasmobranchs which are generalists (Wilga et al. 2007), planktivorous elasmobranchs feed primarily on zooplankton (Sims et al. 2004, Stevens 2007, Nakaya et al. 2008, Couturier et al. 2013) which have an abundance and distribution driven primarily by predictable environmental changes (Richardson 2008). Past studies have found that the aggregations, abundance, behavior, and movement of planktivorous elasmobranchs can be based on predictable food-pulses of zooplankton. In the eastern tropical pacific electronic tagging of whale sharks revealed foraging behavior is associated with high primary productivity (Guzman et al. 2022). Similar phenomena have been documented near La Paz, Mexico, where juvenile whale sharks aggregate to feed on copepod blooms (Clark and Nelson 1997) and at the Belize barrier reef where whale sharks form aggregations which correspond with the spawning aggregations of various species of snapper (Heyman et al. 2001, Graham et al. 2006). Some studies have revealed similar patterns for planktivorous rays: on the Great Barrier Reef manta rays feed in locations and at times of higher zooplankton biomass (Armstrong et al. 2016) and the presence of giant devil rays in the eastern Pacific is influenced by seasonal upwelling events (Lezama-Ochoa et al. 2019).

In addition to food availability, another key factor which influences planktivorous elasmobranch aggregations and behavior is environmental conditions and characteristics. In the western Indian Ocean and off Ningaloo Reef, Australia, the number of whale sharks in pelagic surface waters is strongly correlated with sea surface temperature (SST) (Sequeira et al. 2012). Another study which considered SST, Chlorophyll A, and bathymetry is related to all of these (Sleeman et al. 2010b). In addition, the Southern Oscillation Index and wind shear are linked to abundance and distribution of planktivorous elasmobranchs (Wilson et al. 2001, Sleeman et al. 2010b). Across taxa, lunar cycle has been found to influence species abundance and behavior. Tidal density, new, and full moons increase the aggregations of mantas at Komodo Marine Park (Dewar et al. 2008). Surveys of fishers in Ghana revealed best catches are reported when there is partial or no lunar illumination (Seidu et al. 2021). The largetooth sawfish, a large shark-like ray has diel behaviors which are influenced by lunar illumination levels (Whitty et al. 2017). There is also evidence that foraging behavior is influenced by lunar phase; for example, cape fur seal predation by white sharks is reduced during full moons (Hammerschlag et al. 2006) and white shark sightings occur with the lowest frequency at full moon and the peak at new moon (Weltz et al. 2013). Finally, in some bony-fishes moonlight-related periodicities is used as reliable information for synchronizing the timing of reproductive events (Ikegami et al. 2014).

There is clear potential for environmental variability to influence the abundance and behavior of planktivorous elasmobranchs, and other marine species. However, studies on the efficacy of marine protected areas, assessments of elasmobranch population dynamics, and studies of elasmobranch movement ecology do not always account for environmental factors or simply include one environmental covariate (e.g., temperature or depth) (Goetze and Fullwood 2013, Juhel et al. 2018, Albano et al. 2021, Hammerschlag et al. 2022). However, incorporating a broad array of environmental factors, especially those associated with climate change is essential in making well-supported ecological inferences and generating critical data for management. With respect to planktivores, there are some studies to date which have examined the potential effects of different environmental covariates on planktivorous elasmobranchs, most of these studies are short-term, anecdotal, observational, based on small sizes, or account for limited covariates. Long term datasets are critical to develop an understanding of the ecology and biology of planktivorous elasmobranchs (Stewart et al. 2018, White 2019).

In the present study, we leverage the recent advent of open access remotely sensed environmental data (e.g., open access Chlorophyll A, salinity, and lunar cycle data) to assess trends in the populations of planktivorous elasmobranchs in a 28 year Underwater Visual Census (UVC) dataset. This UVC represents one of the longest underwater censuses of sharks and rays. By using this dataset, we can untangle the many previously suggested relationships between environmental covariates and planktivorous elasmobranchs. The specific aims of this study are threefold. First, we expand on past studies on the same system (White et al. 2015, Osgood et al. 2021) to examine the role of food availability (e.g., primary productivity), acute climate changes (e.g., sea surface temperature, temperature at depth), long term climate trends (e.g., Ocean Niño Index), oceanographic conditions (e.g., current, salinity), and lunar cycle (e.g., lunar phase, lunar distance) on the population trends of planktivorous elasmobranchs. Second, we examine how accounting for environmental covariates shifts predictions about population trends when compared with models which examined population trends over time without accounting for environmental variability. Finally, we examine species-specific differences in the influences of environmental covariates.

## 2. Materials and Methods

### 2.1 Study Site and Underwater Visual Census Protocol

Cocos Island National Park (Isla del Coco; 05°31′08′′, W 87°04′18′′) is a small (23.85 km2) uninhabited island 550 km from mainland Costa Rica, its Marine Protected Area (MPA) was established in 1984, making it the world’s oldest MPA (White et al. 2015) (Figure 1).

**Figure 1:**
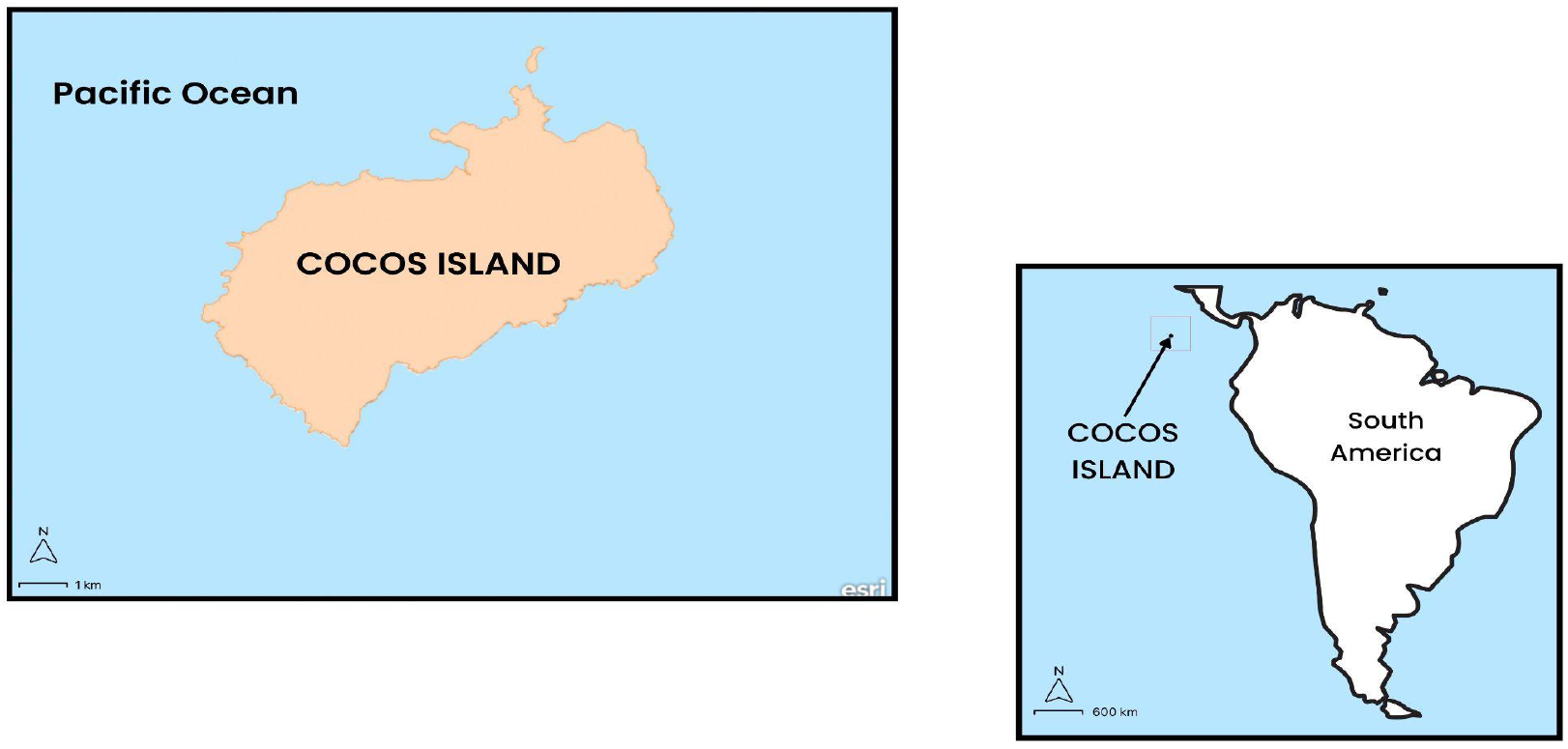
Map of Cocos Island with the location of Cocos in relation to South America.

**Figure 2:**
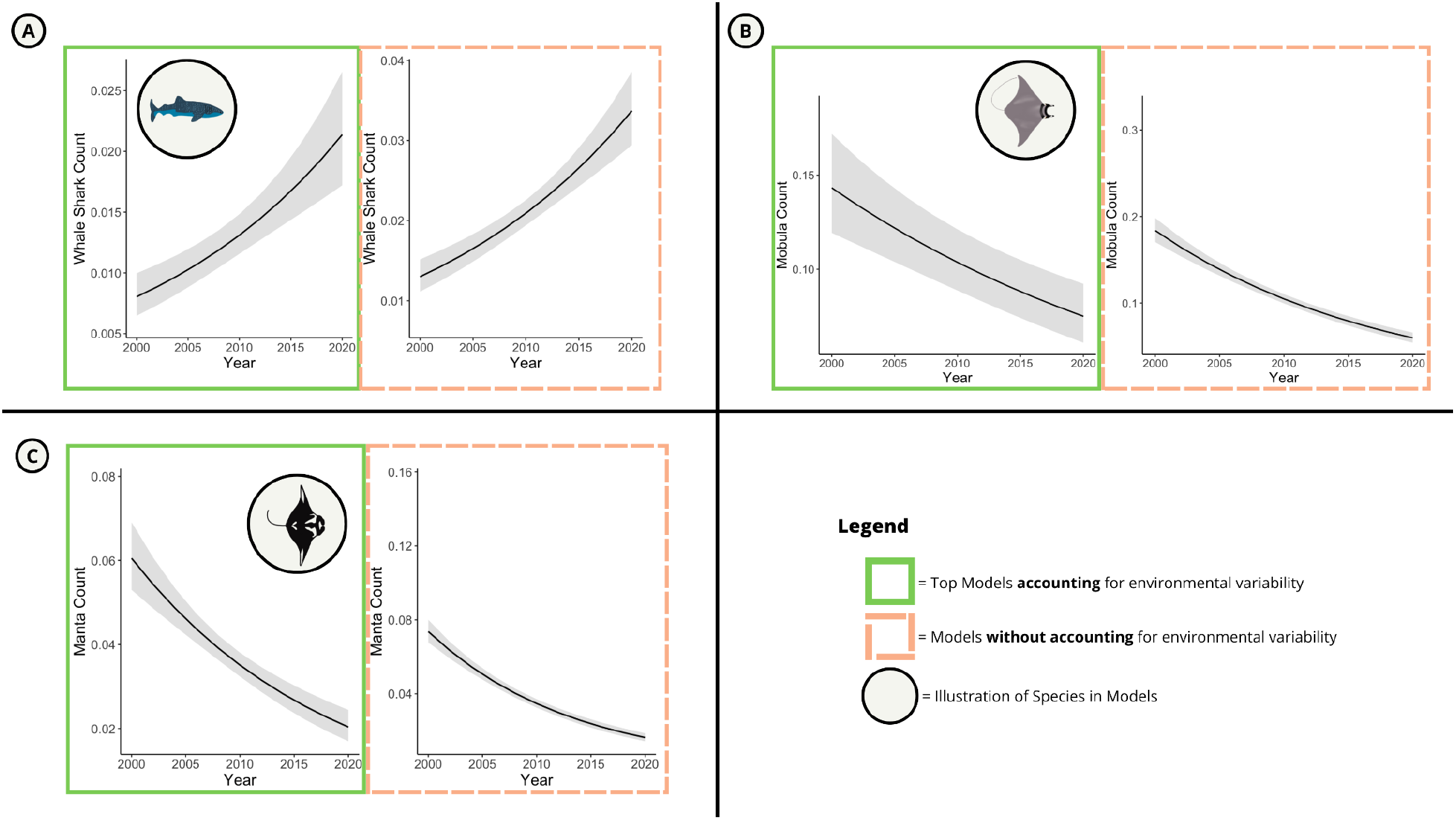
Model comparison for abundances over time when accounting for Ocean Niño Index, temperature at depth, sea surface temperature, salinity, Chlorophyll A, current, visibility, year, seasonality, lunar phase (only mobulas), and lunar distance (only mobulas) [outlined in green] and accounting only for year [outline in orange]. Comparisons are shown for whale sharks (A), mobulas (B), and mantas (C).

**Figure 3:**
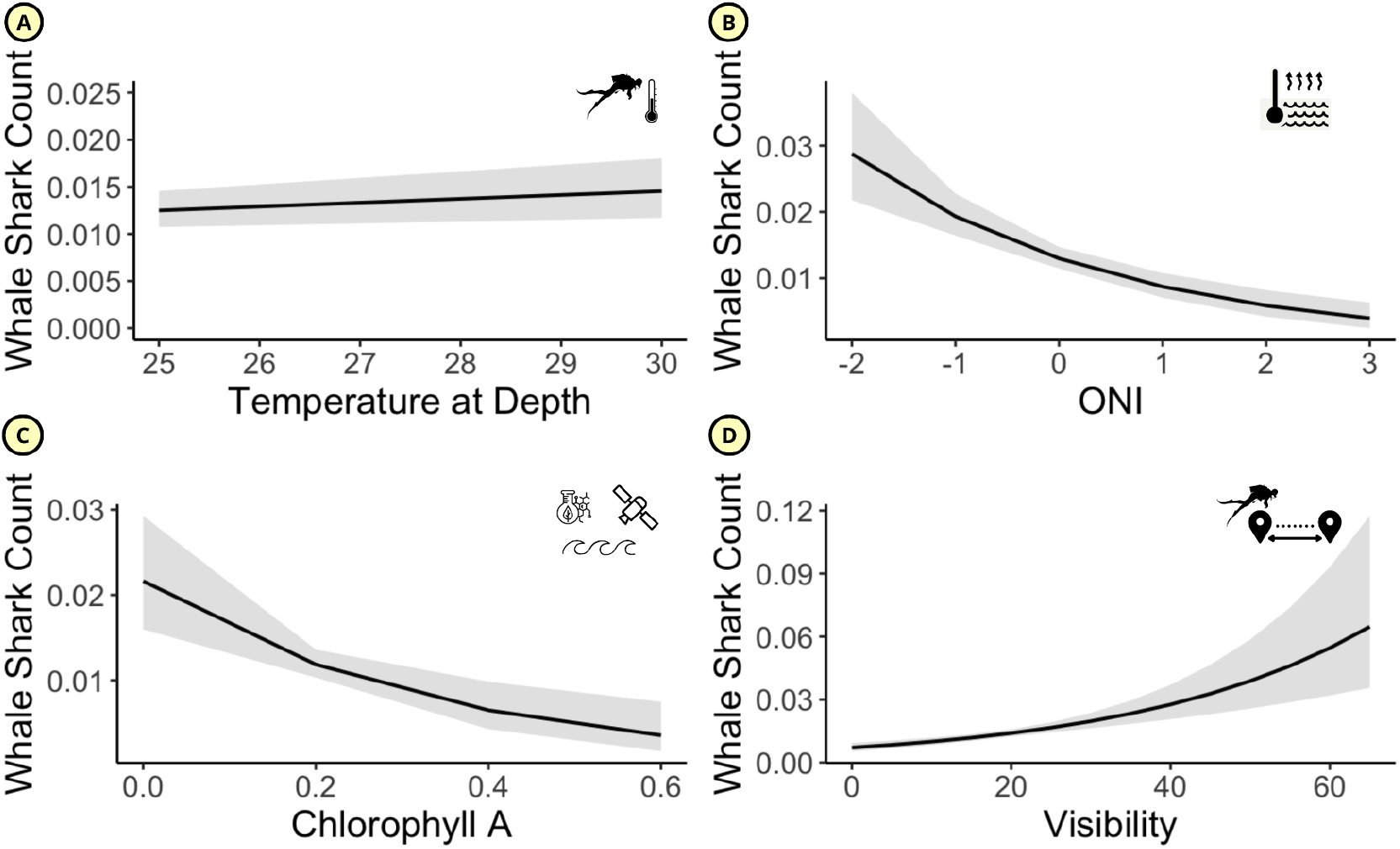
Predicted whale shark count for statistically significant (continuous) covariates (a) temperature at depth, (b) Ocean Niño Index, (c) Chlorophyll A, and (d) water visibility. 95% confidence intervals displayed in gray.

From January 1993 to December 2020 (n = 28 years), experienced diver guides (n = 46) at Undersea Hunter (Figure 1), conducted a total of 35,706 dives at 17 sites around Cocos Islands. This data represents one of the largest underwater visual censuses (UVC) for sharks and rays (White et al. 2015). These dives cannot be considered a scientific UVC because there was no defined field of view. However, protocol was consistent throughout the duration of the study. Dives averaged ∼60 minutes and were led by experienced and trained divemasters. At each site, depth was consistent but overall ranged from 10 to 40 m. Cocos provides exceptional habitat for marine organisms with its location at the nexus of reef and seamount complexes and at the confluence of major ocean currents, Cocos Island is truly a biodiversity hotspot, home to the unique intersection of pelagic and coastal species. Oceanic Islands, like Cocos, provide important habitats in the pelagic environment because they are “borders’’, allowing for reef-associated communities to interact with pelagic species of many different trophic levels (Friedlander et al. 2012). In this study, the dive sites encompassed the range of shallow-water environments and seamount complexes at Cocos Islands.

Upon completion of each dive, divers used a standardized datasheet to record the observed number of several elasmobranch species (*Sphyrna lewini, Triaenodon obesus, Aetobarus narinari, Taeniura meyeni, Mobula spp*., *Manta birostris, Rhincodon typus, Galeocerdo cuvier, Carcharhinus falciformis, Carcharhinus limbatus, Carcharhinus albimarginatus)*.

We transcribed and compiled all data from Undersea Hunter’s 53 divemasters into a single database, applied several filters, and corrected for transcription errors. After this process, 35,706 individual dives conducted by 36 divemasters remained for analysis. In this study, we examined the environmental influences on planktivorous elasmobranchs (*Mobula spp*., *Manta birostris, Rhincodon typus)*. As such, these species are the primary focus of analysis (response variables).

### 2.2 Environmental Data

In addition to the primary UVC data (counts and presence/absence) divemasters recorded several environmental parameters (current, visibility, temperature at depth). We supplemented this survey data with several open-source environmental datasets which included mean monthly sea surface temperature, lunar phase, lunar distance, lunar illumination means, mean monthly salinity, mean monthly chlorophyll A, and Ocean Niño Index (ONI) (Table 1). The Hadley EN4 subsurface salinity objective analysis was used to create a 60+ year of sea surface salinity (at 5 meters depth); we then integrated this data with our original dataset and used it as the salinity covariate. Chlorophyll A data was available beginning in 2002, using the NASA combined-satellite (NASAcombo) time series, a multiple-satellite cross-calibrated chlorophyll product, to create a time series of primary productivity to use as our Chlorophyll A covariate. We selected the Chlorophyll A covariate to serve as a proxy for zooplankton. Chlorophyll A is an indicator of phytoplankton; phytoplankton and zooplankton abundance are often correlated (Ware and Thomson 2005, Richardson 2008). Furthermore, Chlorophyll A has been used as a proxy for zooplankton levels in several other studies on planktivorous marine megafauna (Burtenshaw et al. 2004, Hlista et al. 2009, Sleeman et al. 2010b, Rohner et al. 2018, Harris et al. 2021, Shaw et al. 2021). Chlorophyll A, however, is not a perfect indicator of zooplankton abundance. We also integrated sea surface temperature covariate using a high-resolution blended product based on satellite and in-situ data (HadISST). We obtained lunar data (e.g., lunar distance, lunar phase, and lunar illumination mean) using the package ‘lunar’ (Lazaridis 2014) in R (Core Development Team 2013). Using the same methods as a previous study on the same system, ONI data was obtained from NOAA at their website https://www.cpc.ncep.noaa.gov/data/indices/oni.ascii.txt (Osgood et al. 2021). We adapted the methods from Osgood et al. 2021 and included temperature of dive, mean monthly sea surface temperature, and ONI index. We selected these because they each represent a different type of temperature change: ONI represents the running 3 month mean of SST anomalies in the Niño-3.4 region of the east-central Pacific and correlates with more general oceanographic features of the eastern tropical Pacific, temperature of dive captures the immediate responses of species at depth, and sea surface temperature captures the immediate responses of species at the surface. SST and Temperature at depth were not correlated (rho = 0.3361943), temperature at depth and ONI were not correlated (0.2898731), and SST and ONI were not correlated (rho = 0.3359939).

**Table 1:** all covariates (environmental data) used in models with a description of their respective sources.

| Variable | Range (5th- 95th percentile) | Source | Description | Fixed or Random Effect |
| --- | --- | --- | --- | --- |
| Sea Surface Temperature | 26.17-29.62 | NASA-MODIS<br>https://terra.nasa.gov/data/modis-data | Daily SST | Fixed |
| Ocean Niño Index (ONI) | -1.38-1.31 | NOAA | multivariate El Nino Southern Oscillation index | Fixed |
| Chlorophyll A | 0.108-0.257 | NASACombo time series:<br>https://oceancolor.gsfc.nasa.gov/ | a multiple-satellite cross-calibrated chlorophyll product | Fixed |
| Salinity | 31.47100- 33.42025 | Met Office Hadley Centre observations datasets:<br>https://www.metoffice.gov.uk/hadobs/en4/ | Hadley EN4 subsurface salinity objective analysis 3 | Fixed |
| Lunar Phase | new Moon, waxing crescent, first quarter, waxing gibbous, full Moon, waning gibbous, third quarter and waning crescent | ‘lunar’ package in r:<br>https://cran.r-project.org/web/packages/lunar/lunar.pdf | The category names which correspond to phases of the moon on a given day at Cocos Island. | Fixed |
| Lunar Distance | 56.61736- 63.66912 | ‘lunar’ package in r:<br>https://cran.r-project.org/web/packages/lunar/lunar.pdf | Distance to the moon returned in units of earth radii, or as a 5-level factor variable referring to the moon’s perigee (at about 56 earth radii) and apogee (at about 63.8 earth radii). | Fixed |
| Dive Temperature | 24-29 | Original dataset | Water temperature recorded by dive masters on their personal dive computers at depth | Fixed |
| Current | 0 (none) - 4 (strong) | Original dataset | Estimation of current strength by divemaster | Fixed |
| Visibility | 10-30 | Original dataset | Water visibility (meters) | Fixed |
| Year | 1998-2019 (full range) | Original dataset | Year of study | Fixed |
| Site | 17 unique sites | Original dataset | Identification number of each dive site | Random |
| Divemaster | 36 unique divermasters | Original dataset | Identification number of each divemaster | Random |

### 2.3 Modeling Environmental Influences on Elasmobranchs

We modeled the counts of each focal species using a hierarchical generalized linear mixed model (GLMM) framework. We selected these mixed effect models to implement random effects to account for observations made by the same divemaster or at the same dive site. To account for seasonality, we included the sin and cosine functions of Julian date as explanatory variables (Baum and Blanchard 2010). We adapted each model to the data type and distribution of the species. For all three species included in our models, count data were recorded; however, these species were rarely observed. Because of the low frequency of observation, we employed zero-inflated mixed models with negative binomial distribution (ZINB). All models were implemented in the ‘glmmTMB’ package (Magnusson et al. 2017) in R (Core Development Team 2013).

For model selection, rather than using a “drop-one” approach for AIC, we choose to select a series of models which each have a biologically sensible (see supplementals) and plausible interpretation (Arnold 2010). We used the ‘AICcmodavg’ (Mazerolle 2020) package in R (Core Development Team 2013) to select top models from our series of biologically sensible models (see supplementals). If several models fell within 2 AIC of the best model, we used a model averaging approach (Zuur et al. 2009) to generate parameter estimates. We employed this model averaging approach with the ‘model.avg’ function in the ‘MuMIn’ package (Bartoń 2015) in R (Core Development Team 2013).

## 3. Results

### 3.1 Model Comparison

We found significant changes in model interpretations when we accounted for environmental covariates as opposed to only including year as a covariate. For whale sharks, the model which did not account for environmental variability and the model which included environmental covariates estimated similar trends (i.e., population increases each year), but underestimated the increases in populations. Our top model estimated a 6% increase in whale shark relative-abundances each year (p<0.001), whereas our model which did not account for environmental variability estimated a 5% increase in whale shark relative-abundances each year (p<0.001). For mobula rays, failure to account for environmental covariates leads to underestimate of population increases. Our model average of our top two models estimated a 5% decrease in mobula ray relative abundance each year (p<0.001); however, our model which did not account for environmental variability estimated a lesser 3% decrease in relative abundance of mobulas each year (p<0.001). Finally, accounting for environmental variability our top model estimated a 5% decrease (p<0.001) in the relative abundance of manta rays each year, while our model which did not account for environmental variability estimated a larger 6% decrease in relative abundance of manta rays each year (p<0.001).

### 3.2 Whale Sharks

Whale sharks were observed on 2.11% of dives. The top model for whale sharks was the global model without lunar covariates. We estimated that over the last two decades, there was a significant increase in the abundance of whale sharks; specifically, we modeled that each year there is a 6% increase in the abundance of whale sharks (p<0.001). We modeled no significant effect of SST on whale shark relative abundance; however, a 1°C in temperature at depth yielded a 6% increase in the abundance of whale sharks (p=0.031). Additionally, long term climate trends appeared to influence whale shark abundance, with a 1 unit increase in ONI yielded a 30% decrease in the abundance of whale sharks (p<0.001). We also modeled that a 0.10 *mg*/*m*^3^increase in surface Chlorophyll A yielded a 26% decrease in the abundance of whale sharks (p<0.001). Finally, we modeled that with a 1 m increase in visibility there is a 3% increase in the abundance of whale sharks (p=0.001).

### 3.4 Mobula Rays

Mobula rays were observed on 11.9% of dives. For mobula rays, two models fell within 2 AIC of each other (Table 2), the following results are based on a model averaging approach of our global model without lunar factors and our global model without salinity. We found that over the last two decades, there was a statistically significant decrease in mobula ray abundance. Specifically, we modeled a 5% decrease in the abundance of mobulas each year (p<0.001). Increases in temperature at the surface and at depth yielded significant decreases in the abundances of mobula rays; specifically, we modeled that a 1°C increase in SST yielded a 22% decrease in the abundance of mobulas (p<0.001), and a 1°C increase in temperature at depth yielded a 4% decrease in the abundance of mobulas (p=0.049). We also found that climate trends influenced the abundance of mobulas, with a 1 unit increase in ONI yielding a 17% increase in the abundance of mobulas (p=0.002). While primary productivity did not have a significant effect on the abundance of mobula rays, salinity did have a significant effect on mobulas. We modeled that a 1 unit increase in surface PSU leads to a 16% increase in the abundance of mobula rays (p=0.048). Two lunar phases appeared to have significant effects on mobula ray abundance; we modeled that during the last quarter there is a 34% increase in mobulas relative to the first quarter (p=0.016) and during the new moon there is a 42% increase relative to the first quarter (p=0.004). Finally, we found that increases in current and visibility increased the abundance of mobulas. A 1-meter increase in visibility yielded a 4% increase in the abundance of mobulas (p<0.001) and a 1 unit increase in current strength yielded a 12% increase in the abundance of mobulas (p<0.001).

**Table 2:** Top model and model type for each species included in this study; *is an indicator of significance (P ≤ 0.05); all models also include sin and cosine of Julian Date to account for seasonality; Abbreviations: GLMM, generalized linear mixed model; ZI, zero-inflated. Each model also includes the effects of the site and divemaster.

| **Species** | **Top Model(s)** | **Model Type** |
| --- | --- | --- |
| Whale Sharks | Ocean Nino Index* + Temperature at Depth* + Sea Surface Temperature + Salinity + Chlorophyll A* + Current + Visibility* + Year* | GLMM, ZINB |
| Mobula Rays | A. Ocean Nino Index* + Temperature at Depth* + Sea Surface Temperature* + Salinity* + Chlorophyll A + Current* + Visibility* + Year *<br>B. Ocean Nino Index* + Temperature at Depth* + Sea Surface Temperature* + Chlorophyll A + Lunar Distance + Lunar Phase* + Current *+ Visibility* + Year* | GLMM, ZINB |
| Manta Rays | Ocean Nino Index + Temperature at Depth* + Sea Surface Temperature* + Salinity + Chlorophyll A + Current + Visibility* + Year* | GLMM, ZINB |

### 3.5 Manta Rays

Manta rays were observed on 4.21% of dives. Like with whale sharks, the top model for manta rays was the global model without lunar-related parameters (Table 2). Our top model estimated that over the last two decades, manta ray abundance decreased each year by 5% (p<0.001). Our models found no statistically significant effect of primary productivity, salinity, or current on manta rays. Additionally, mantas did not appear to be affected significantly by ENSO. However, mantas did appear to be sensitive to acute temperature changes; specifically, our models estimated a 1°C increase in temperature at depth leads to a 6% decrease in the abundance of mantas (p=0.026). More impactful than temperature at depth was SST, where a 1°C increase yields a 16% decrease in the abundance of mantas (p=0.014). Finally, we modeled that a 1-meter increase in visibility yields a 2% increase in the abundance of mantas (p<0.001).

## 4. Discussion

### 4.1 Population Trends and Model Comparisons

We observed significant decreases in the abundances of both mantas and mobulas over time. Rays which exhibit aggregations are highly vulnerable to exploitation; the aggregations of mantas make them particularly susceptible to becoming bycatch in purse seine and longline fisheries (Duffy and Griffiths 2017). Giant manta rays are considered easy to target because of their “large size, slow swimming speed, tendency to aggregate, predictable habitat use, and lack of human avoidance” (Marshall et al. 2020). Retainment during incidental catches of manta rays is common because of their high trade value (Croll et al. 2016, Lawson et al. 2017, Marshall et al. 2020), and even when released alive, they are often injured from being captured, and suffer high post-release mortality (Tremblay-Boyer and Brouwer 2016).

We observed an increase in the abundance of whale sharks over time. We believe a possible explanation for this is the decrease in abundance of manta and mobula rays, since they occupy a similar ecological niche, as pelagic filter-feeders in tropical seas, where plankton is scarce (Couturier et al. 2013). Similar niche partitioning of elasmobranchs has been observed in other elasmobranchs; for example, in South Africa, there is empirical evidence for behavioral shifts of sevengill sharks following the decline and disappearance of white sharks from a foraging site (Hammerschlag et al. 2019). Additionally, in Hawaii, data from dietary analysis revealed that interspecific competition highly influences the distribution of carcharhinid sharks (Papastamatiou et al. 2006). Our results may be an example of one of the most basic ecological principles, competition.

One of the goals of this study was to examine what happens when models do not account for environmental variability and how this affects interpretations of model outputs. In our models for all three species, there were significant differences in the estimated year parameter when we accounted for environmental variability. For manta rays and whale sharks, when we did not account for environmental variability in our models, we overestimated population changes each year. For whale sharks, we estimated a larger increase in populations and for mantas we estimated a larger decrease in populations. For mobula rays we underestimated population decreases when we failed to account for environmental variability, this is likely because mobulas were highly influenced by a variety of environmental covariates (see figure 4). Since studies like these are often used to inform management strategies and IUCN statuses, discrepancies in estimations can be problematic. We show the necessity to account for environmental variation when examining population trends.

**Figure 4:**
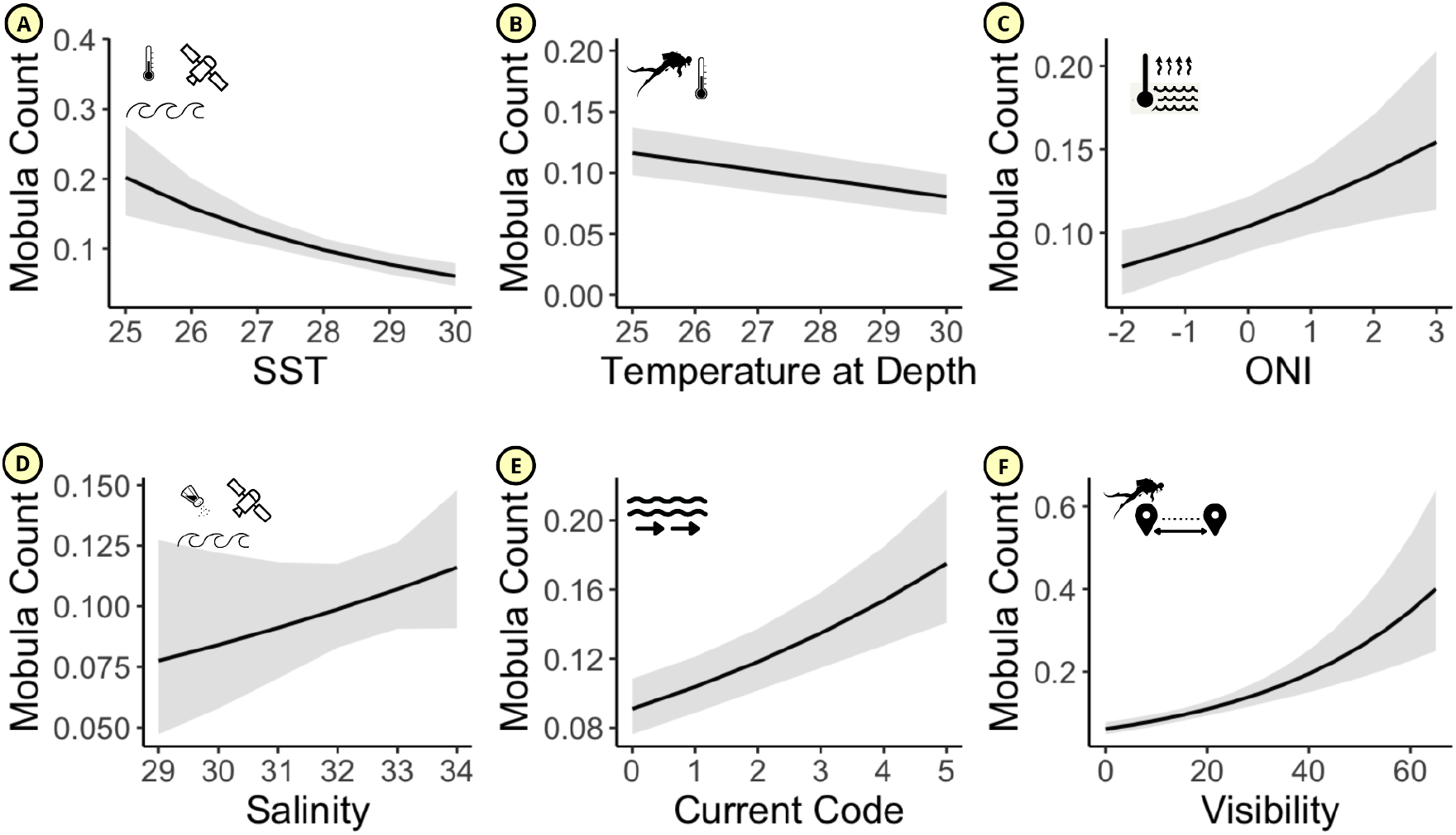
Predicted mobula count for statistically significant (continuous) covariates (a) sea surface temperature (SST), (b) temperature at depth, (c) Ocean Niño Index, (d) salinity, (e) current, and (f) water visibility. 95% confidence intervals displayed in gray.

**Figure 5:**
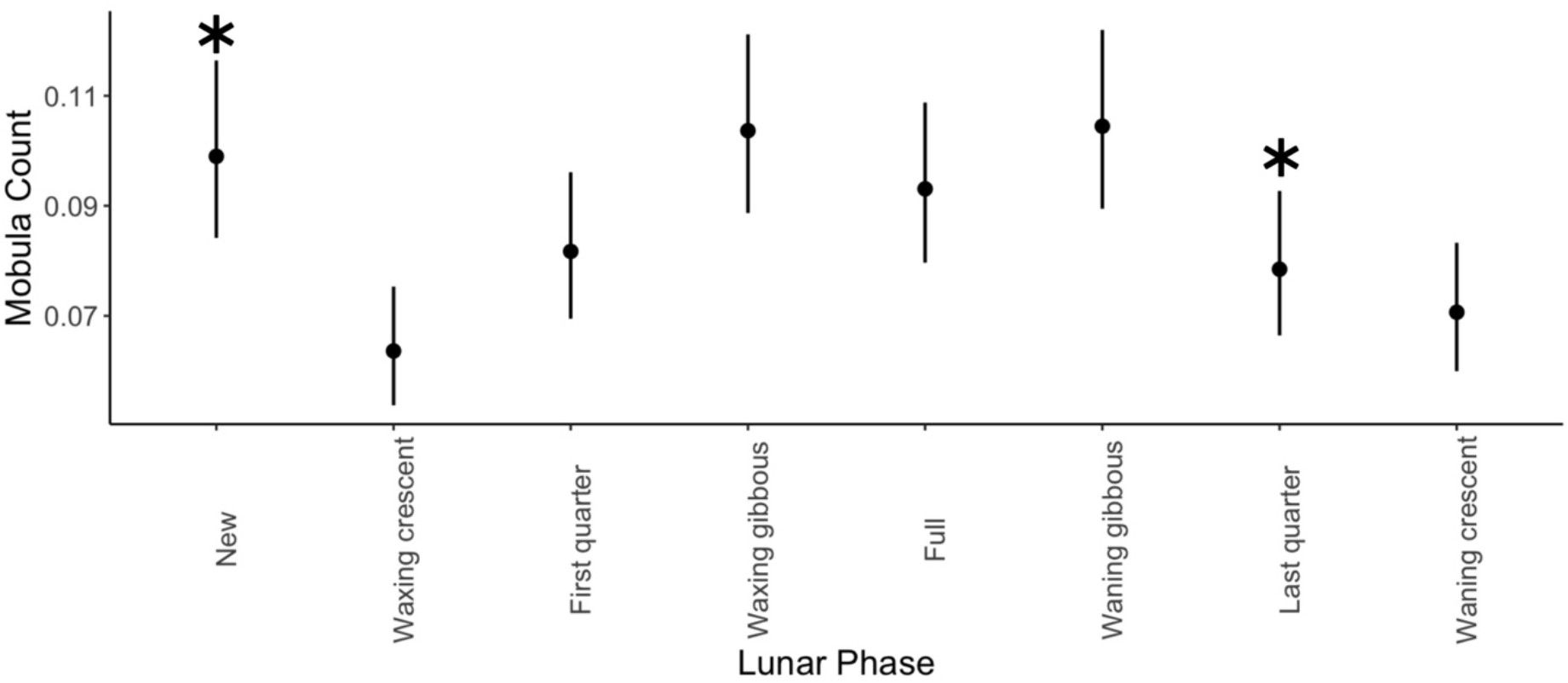
Effects of the categorical variable on mobula rays lunar phases with statistical significance relative to the first quarter p < 0.05 indicated by a *.

**Figure 6:**
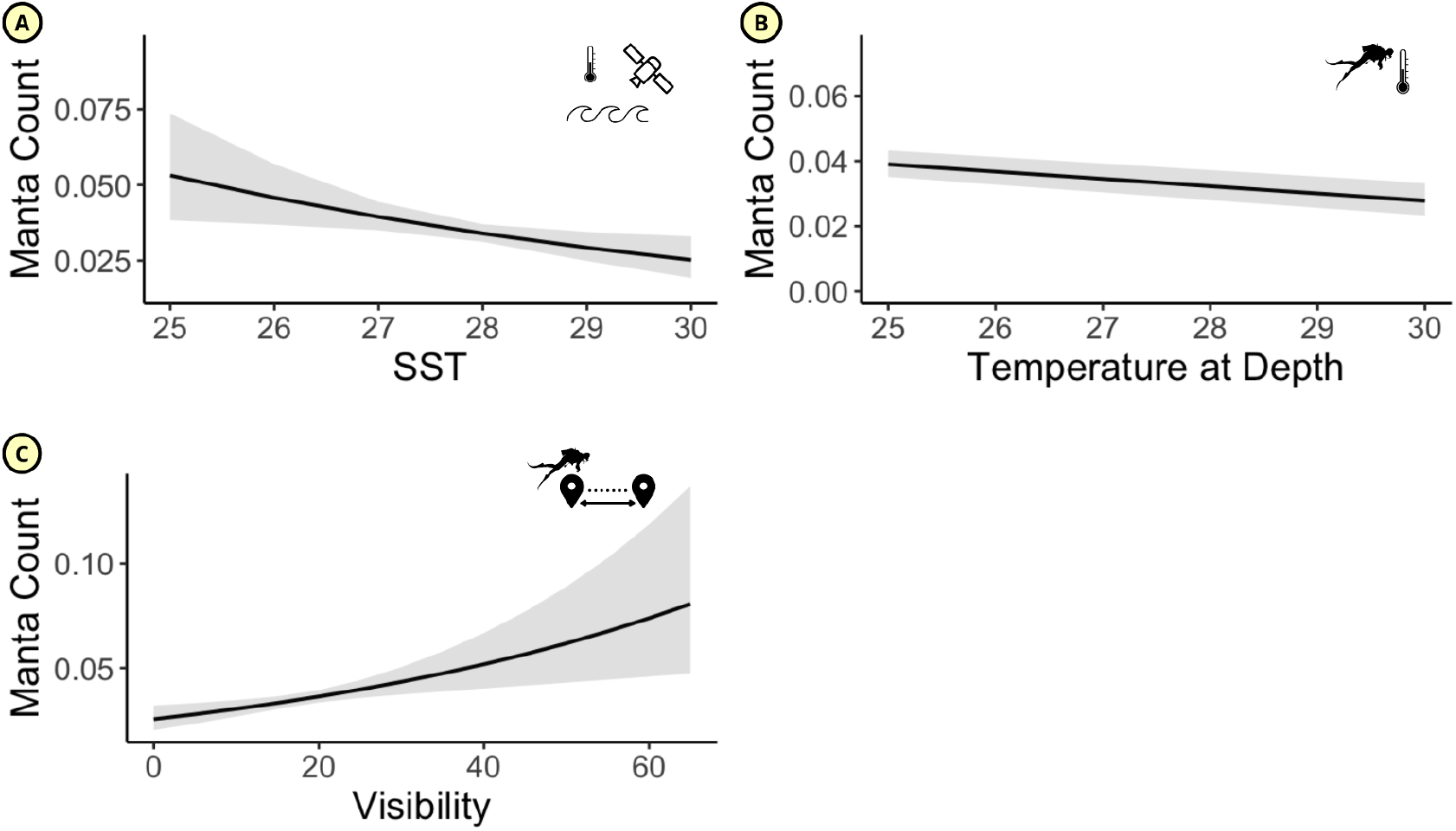
Predicted manta count for statistically significant (continuous) covariates (a) sea surface temperature (SST), (b) temperature at depth, and (c) water visibility. 95% confidence intervals displayed in gray.

### 4.2 Temperature and Climatic Conditions

Our models showed no significant relationship between SST and whale shark abundance; however, we did model a 39% decrease in the abundance of whale sharks with a 1 unit increase in ONI and a 6% increase in abundance of whale sharks for a 1°C increase in temperature at depth. The results of our study support the findings of several past studies which found that El Niño and La Niña influence the abundance of whale sharks (Wilson et al. 2001, Sleeman et al. 2010a). It is thought that whale shark movements are related to ocean currents, which are fundamentally shifted during La Niña and El Niño years. One potential explanation for this is the physiology of whale sharks, who like many other species of large fish, are known to have a wide thermal tolerance– a 2020 study suggested the large body size of whale sharks prevents decrease in temperature during deep excursions without high metabolic costs to maintain body heat (Nakamura et al. 2020).

For mobula rays and manta rays, we found that increases in temperature yielded significant decreases in their abundance. Mobulas were influenced by SST, with a 1°C increase in SST leading to a 12% decrease in abundance. Temperature at depth also played a small but significant role in the abundance of mobula rays, with a 1°C increase in temperature at depth yielding a 4% decrease in the abundance of mobulas. Manta rays were influenced by both temperature at depth and SST, with a 1 °C increase in SST yielding a 13% decrease in abundance and a 1°C increase in temperature at depth yielding a 6% decrease in abundance. One of the key takeaways of Osgood et al. 2021 was that mobula rays (mantas and mobulas combined) showed little response to ONI. Unlike Osgood et al. 2021, we choose to keep the manta ray category and mobula ray category separate, with mantas being *Manta birostris* and mobulas being all other members of the *Mobula* genus surveyed. Our results for mantas were similar to that of Osgood, finding no significant relationship between mantas and ONI. However, ONI did play an important role in mobula abundances, with a 1 unit increase in ONI yielding a 17% increase in mobula abundance. While in this study we show that acute temperature changes, specifically warming, lead to decreases in abundance of rays, other studies have shown that ocean warming can exert a variety of effects on elasmobranchs. For example, warming can shift freeze responses in elasmobranchs leading to greater vulnerability to predation (Ripley et al. 2021). Another study found ocean warming can affect the development of sharks, with patterns which are characteristic of species developing incorrectly when individual sharks are reared at high temperatures (Gervais et al. 2016).

### 4.3 Primary Productivity

To better understand how elasmobranch abundance is related to physical transport mechanisms and primary productivity, we decided to include Chlorophyll A concentration. We hypothesized that high primary productivity would increase abundance of planktivores because they are known to follow physical and biological oceanographic cues to locate successful foraging grounds (Nelson and Eckert 2007). Our results, however, were in contrast with this hypothesis. We found that manta ray and mobula ray abundance was not significantly affected by primary productivity. For whale sharks, increased primary productivity decreased abundance. Our models found that for 0.10 *mg*/*m*^3^ increase in Chlorophyll A there was a 26% decrease in the abundance of whale sharks.

Upon examination of this, there are some plausible explanations for this seemingly odd phenomenon of planktivores avoiding high primary productivity. Throughout the duration of this study, divemasters noted that unlike at other known whale shark aggregations, the whale sharks at Cocos did not appear to be feeding. It is possible that the whale sharks surveyed in this study were not practicing surface feeding, but rather exhibit deep-dive foraging behavior, which the species is known to exhibit in other locations (Graham et al. 2006). Tracked whale sharks in the Sea of Cortez spend long periods at depth and short sporadic times near the surface (Eckert and Stewart 2001). Another study which found a lack of strong correlation between whale shark movement patterns and measured Chlorophyll A suggested that Chlorophyll A is a poor proxy for zooplankton biomass (Sleeman et al. 2010b). Chlorophyll A being a poor proxy for zooplankton abundance is supported by literature: for example, a study which found that systems with larger amounts of macrozooplankton relative to microzooplankton have less Chlorophyll A relative to total phosphorus, suggesting that large zooplankton (the preferred prey of planktivores) may actually reduce Chlorophyll A levels (Pace 1984). Another study found no major changes in zooplankton abundance and community structure with increased concentrations of Chlorophyll A (Canfield and Watkins 1984). Another possibility is that whale sharks are responding to changes in zooplankton community composition (i.e., specific species present), rather than the concentrations of species. Neither White et al. 2015 or Osgood et al. 2021 included Chlorophyll A in their models; however, in our models we included this parameter, and it was one of the most influential covariates in our whale shark models.

There is, however, a notable limitation of our data obtained through satellite measurements: data only indicates levels of primary productivity at the sea surface. Species of filter feeders are known to exhibit broad-scale habitat use across surface and pelagic zones. Basking sharks *Cetorhinus maximus* dive to forage on zooplankton communities in the mesopelagic and epipelagic zones (Sims et al. 2003). A study which investigated whale shark diving patterns on the Mesoamerican barrier reef suggested that whale shark diving patterns (including deep dives) may be influenced by a seasonally available food source (Graham et al. 2006). Keeping in mind the known deep-diving behavior of planktivores, in order to truly understand the role of primary productivity and Chlorophyll A concentrations in the behavior of planktivores, measurements of Chlorophyll A and plankton abundance beyond surface measurements are necessary. We choose to include Chlorophyll A in our models because for this location it was the best available proxy for zooplankton abundance; however, the relationship between Chlorophyll A and zooplankton abundance is a complex non-linear relationship which varies on the species-level.

### 4.4 Salinity

Another commonly assessed oceanographic parameter is salinity. Salinity had a large effect on the abundance of mobula rays; we found that for one unit increase in surface PSU there is a 22% increase in mobula rays. Salinity did not have a statistically significant effect on the abundances of whale sharks or manta rays. A past study on whale shark behavior found that dives were not associated with hydrographic features (salinity and temperature) and cited food availability as the primary reason for varying behaviors (Gunn et al. 1999). One of the few studies on the spinetail devil ray, *Mobula mobular*, found that their presence is influenced by a variety of environmental variables, including salinity (Lezama-Ochoa et al. 2019).

Nevertheless, other studies on marine megafauna, specifically elasmobranchs have found that salinity can play an important role in physiology, metabolism, and behavior. Physiologically, elasmobranchs can adapt to low salinities; a study on captive sharks found coordinated molecular responses to low salinity in the rectal glands and gills (Dowd et al. 2010). Similarly, a study on the habitat partitioning of bull sharks *Carcharhinus leucas* found that juvenile bull sharks may have specific salinity preferences which affect their habitat use (Simpfendorfer et al. 2005). The results of our study may point to a similar phenomenon for mobula rays; they may prefer waters with higher salinities. However, for manta rays and whale sharks, factors other than salinity have a greater effect on their abundance and behavior. Notably, there is limited literature on the effects of salinity on whale sharks, as many studies of whale sharks are results of field observations or deployment of electronic tags. With this in mind, we suggest that future studies, especially those which employ electronic tagging, integrate salinity data as a covariate to further examine the role of salinity in the habitat use and behavior of sharks.

### 4.5 Lunar Factors

Lunar factors are known to impact reproduction (Szmant-Froelich et al. 1985, Perea et al. 2022), migration (Somers and Stechey 1986, Sleeman et al. 2010a, Norevik et al. 2019), behavior (Naylor 2001, Mestre et al. 2019), and physiology (Portugal et al. 2019) across different species and ecosystems. For planktivorous elasmobranchs at Cocos, influence of lunar factors ranged from no influence to significant or large influence. On the larger-bodied manta ray and whale sharks, lunar factors were not included in top models. This is in contrast to some past work at the Belize Barrier Reef, whale sharks for seasonal aggregations around predictable spawning aggregations of various species of snapper (Heyman et al. 2001, Graham et al. 2006) and at Ningaloo Reef aggregations of whale sharks are formed around the timing of known coral spawning events (Gunn et al. 1999).

We found that for mobula rays there is a significant increase in abundance during new moons and last quarters, relative to the first quarter. One potential explanation for this is the increase in lunar illumination during and leading up to full moons creates a consistent pattern of increased vulnerability to predation of the smaller-bodied mobula rays. Therefore, their abundance increases under circumstances with lower lunar illumination (i.e., last quarter and new moon). Additionally, the literature on other species of elasmobranchs have suggests that circadian rhythm may regulate diving and metabolic patterns (Nelson and Johnson 1970, Nixon and Gruber 1988). The significant role of lunar factors on mobulas may be due to the known role of lunar cycle on fish and coral spawning and the known influence of these events on the foraging behavior of planktivorous elasmobranchs. However, the fish and coral communities at Cocos Island are relatively under-studied, making it difficult to confirm this.

We hypothesize that a potential reason for the increase in the number of encountered rays when lunar illumination decreases is anti-predatory behavior. In areas where lunar illumination is high it is possible that they move away from foraging locations where increased light at the surface would leave them vulnerable to predation by larger sharks (i.e., scalloped hammerheads, tigers, and other carcharhinid species). Other studies have supported that lunar phase impacts movement and foraging behavior; for example, a study on the movements of gray reef sharks found that the mean depth inhabited increased throughout the lunar cycle (Vianna et al. 2013). If our hypothesis that lunar cycle influences behavior by eliciting an anti-predatory response in smaller-bodied mobula rays under increased light conditions is correct, it is viable that whale sharks and larger-bodied mantas would not exhibit the same antipredatory response since they are both (at this study location) too large to be preyed on by the medium to large-bodied carcharnids.

### 4.6 Limitations and Future Work

One of the major limitations of this study was that Chlorophyll A is not the best indicator of zooplankton abundance, as primary productivity levels and zooplankton abundance are not the same. In the future, especially for studies on planktivorous elasmobranchs, we suggest additional methods to assess levels of zooplankton abundances. For example, a 2019 study developed a satellite-derived proxy of mesozooplankton (Druon et al. 2019). In the future, we could include a covariate like this one in models to indicate food availability for planktivores. Additionally, to truly understand how zooplankton abundances affect planktivores, surveys of zooplankton at the species-specific level in conjunction with observation of planktivores (e.g., zooplankton samples in conjunction with all dive surveys) are necessary. In the future, water samples at known foraging sites could be DNA sequenced to examine assemblages, providing accurate information on abundance and diversity of invertebrates within the water column (Hajibabaei et al. 2011). Likewise, plankton tows or continuous plankton recorders are employed at other locations to monitor plankton assemblages and have potential for use in studies on planktivorous elasmobranchs (Head et al. 2022).

Other studies on planktivores have additional available environmental covariates. for instance, a study which examined the environmental characteristics associated with the presence of the Spinetail devil ray *Mobula mobular* identified chlorophyll and sea surface height as the main predictors of devil ray presence/absence (Lezama-Ochoa et al. 2019). Lezama-Ochoa et al. 2019 included sea surface height values because of their relation to upwelling systems; thus, in the systems where this data is obtainable or monitorable, sea surface height should be included as a predictor.

Although we had high statistical power in our study (White 2019), there could still be potential issues while using community science data. Diver surveys do not have a consistent field of view or length of dive. However, the protocols were the same over 28 years and with a small group of divemasters responsible for all of the data collection. Nevertheless, there could be sampling bias in terms of which sites are visited over time (Fournier et al. 2019; White & Bahlai 2021) Thus, future work could couple standardized surveys with this divemaster-collected data for a more complete picture of populations trends in the area. We suggest that future studies should include Baited Remote Underwater Video Surveys and examine movement of animals via telemetry.

## 5. Conclusion

Our findings highlight that it is critical, when evaluating the efficacy of marine protection, to also consider the potential effects of environmental variability. Many studies evaluate, and often claim success in Marine Protected Areas; however, their work does not account for environmental variability. For example, a study done in South Africa suggested relative abundances of sharks are higher inside of a marine reserve when compared with abundances outside of a marine reserve; however, this study only included depth as a proxy for temperature in their models, but did not have any other environmental covariates (i.e., primary productivity, salinity, tides) (Albano et al. 2021). Similarly, a recent study on white sharks in the Mediterranean Sea analyzed encounter data using generalized additive models and information on human population abundance as a proxy of observation effort; however, they did not account for environmental variation within their models, although they suggest temperature and productivity for potential reasons for fluctuations in abundances (Moro et al. 2020). Nevertheless, other studies have successfully examined shark protection and conservation, while also accounting for environmental variability. A 2020 study used a set of environmental predictors, similar to those which we included in our study to estimate distribution of several species of sharks and propose expansion of protected areas, they suggested that because of decreases in suitable habitat, climate change scenarios should be included as part of shark management strategies (Birkmanis et al. 2020).

In this study, we show that environmental parameters can have significant effects on the abundance and behavior of planktivores. If we fail to include environmental data in models, we could infer increases or decreases in species abundances are the result of policy successes or failures, when the true cause of the shift in species abundances is environmental change. The long-term time series employed in this study provided us with a unique opportunity to explore environmental variability and species trends within an isolated protected area over many years. Additionally, we agree with the conclusions of past studies which suggest that in the future researchers could seek to understand how environmental change affects species interactions in order to predict emergent ecological changes (Kindinger et al. 2022).

## Data Availability Statement

All code for models and data integration is available at https://github.com/juliasaltzman1/Planktivores. The dataset used for this study is property of Undersea Hunter, and is used with their permission, therefore the data is not available online.

## Acknowledgements

The data in this study is property of UnderSea Hunter. We thank all the divemasters and crew members who have worked for the Undersea Hunter dive company over the past 2 decades. This work was improved with feedback from Dr. Elizabeth Harvey, Dr. Catherine Macdonald, Dr. Nathan Furey, Dr. Mario Espinoza, Dr. Alex Hearn, Dr. Remington Moll, Dr. Julia Wester, Dr. Geoffrey Ogood, Dr. Julia Baum, Sophie Wulfing, Ana Silverio, and Wilton Burns. Travel to present this work was made possible by the UNH School of Marine Science and Ocean Engineering Travel Fund and UNH Biological Sciences Travel Fund.

## Supporting Information

Model outputs are available in Supplemental Tables 1-3. Residual plots for each species are available in Supplemental Figures 1-6. Tables with ranking of the top models for each species are in Supplemental Tables 4-6. All models used for AIC along with their descriptions are in Supplement Table 7.

