## Supplemental Information for "Determining the Role of Environmental Covariates on Planktivorous Elasmobranch Population Trends within an Isolated Marine Protected Area"

**Saltzman and White 2022 Supplementals**

***Top Model Outputs***

**Table 1:** whale shark top model output from global model without lunar

| **PREDICTORS** | **INCIDENT RATE RATIOS** | **CONFIDENCE INTERVALS** | **P-VALUE** |
| --- | --- | --- | --- |
| (Intercept) | 0 | 0.00 – 0.00 | <0.001 |
| SST | 0.91 | 0.78 – 1.06 | 0.232 |
| Temperature | 1.06 | 1.01 – 1.12 | 0.031 |
| SALINITY | 0.86 | 0.70 – 1.06 | 0.153 |
| CHLA | 0.05 | 0.01 – 0.24 | <0.001 |
| ONI | 0.7 | 0.60 – 0.80 | <0.001 |
| CurrentCode | 0.96 | 0.90 – 1.03 | 0.299 |
| Visibility | 1.03 | 1.01 – 1.04 | 0.001 |
| SIN TIME | 0.68 | 0.56 – 0.82 | <0.001 |
| COS TIME | 0.27 | 0.23 – 0.32 | <0.001 |
| Year y | 1.06 | 1.04 – 1.08 | <0.001 |
| (Intercept) | 2.01 | 1.49 – 3.38 |  |
| **Zero-Inflated Model** | |  |  |
| (Intercept) | 0 | 0.00 – Inf | 0.994 |
| **Random Effects** | |  |  |
| σ^2^ | 5.36 |  |  |
| τ_00_ _DiverCode_ | 0.03 |  |  |
| τ_00_ _SiteCode_ | 0.81 |  |  |
| ICC | 0.13 |  |  |
| N _DiverCode_ | 45 |  |  |
| N _SiteCode_ | 17 |  |  |
| Observations | 29804 |  |  |
| Marginal R^2^ / Conditional R^2^ | 0.139 / 0.255 |  |  |

**Table 2:** manta ray top model output from global model without lunar

| **PREDICTORS** | **INCIDENT RATE RATIOS** | | **CONFIDENCE INTERVALS** | **P-VALUE** |
| --- | --- | --- | --- | --- |
| Intercept) | $1.5\times{10}^{28}$ | | $1.2\times{10}^{27}-1.9\times{10}^{29}$ | <0.001 |
| SST | 0.87 | | 0.78 – 0.98 | 0.026 |
| Temperature | 0.94 | | 0.90 – 0.99 | 0.014 |
| SALINITY | 1.04 | | 0.88 – 1.23 | 0.651 |
| CHLA | 1.24 | | 0.33 – 4.59 | 0.749 |
| ONI | 1.04 | | 0.93 – 1.17 | 0.515 |
| CurrentCode | 0.96 | | 0.90 – 1.03 | 0.293 |
| Visibility | 1.02 | | 1.01 – 1.04 | <0.001 |
| SIN TIME | 1.08 | | 0.93 – 1.26 | 0.317 |
| COS TIME | 0.92 | | 0.82 – 1.02 | 0.118 |
| Year y | 0.95 | | 0.93 – 0.97 | <0.001 |
| (Intercept) | 1.12 | | 1.10 – 1.15 |  |
| **Zero-Inflated Model** | | |  |  |
| (Intercept) | 0 | | 0.00 – Inf | 0.99 |
| **Random Effects** | |  | |  |
| σ^2^ | 3.62 |  | |  |
| τ_00_ _DiverCode_ | 0.05 |  | |  |
| τ_00_ _SiteCode_ | 0.16 |  | |  |
| ICC | 0.05 |  | |  |
| N _DiverCode_ | 45 |  | |  |
| N _SiteCode_ | 17 |  | |  |
| Observations | 29805 |  | |  |
| Marginal R^2^ / Conditional R^2^ | 0.042 / 0.094 |  | |  |

**Table 3:** mobula ray top model outputs; model average between the global model without salinity and the global model without lunar

| **PREDICTORS** | **INCIDENT RATE RATIOS** | **CONFIDENCE INTERVALS** | **P-VALUE** |
| --- | --- | --- | --- |
| cond((Int)) | $1.4\times{10}^{48}$ | $5.6\times{10}^{5}-3.6\times{10}^{70}$ | <0.001 |
| cond(SST) | 0.78 | 0.70 – 0.88 | <0.001 |
| cond(Temperature) | 0.96 | 0.91 – 1.00 | 0.049 |
| cond(SALINITY) | 1.16 | 1.00 – 1.34 | 0.048 |
| cond(CHLA) | 1.45 | 0.39 – 5.37 | 0.574 |
| cond(ONI) | 1.17 | 1.06 – 1.30 | 0.002 |
| cond(CurrentCode) | 1.12 | 1.05 – 1.19 | <0.001 |
| cond(Visibility) | 1.04 | 1.02 – 1.05 | <0.001 |
| cond(SIN TIME) | 1.71 | 1.50 – 1.94 | <0.001 |
| cond(COS TIME) | 0.87 | 0.79 – 0.97 | 0.01 |
| cond(Year y) | 0.95 | 0.92 – 0.97 | <0.001 |
| zi((Int)) | 0 | 0.00 – Inf | 0.987 |
| cond(LunarDistance) | 0.98 | 0.95 – 1.00 | 0.078 |
| cond(LunarPhase8Full) | 0.84 | 0.66 – 1.08 | 0.176 |
| cond(LunarPhase8Last | 1.34 | 1.06 – 1.69 | 0.016 |
| quarter) |  |  |  |
| cond(LunarPhase8New) | 1.42 | 1.12 – 1.80 | 0.004 |
| cond(LunarPhase8Waning | 1.1 | 0.86 – 1.40 | 0.436 |
| crescent) |  |  |  |
| cond(LunarPhase8Waning | 0.97 | 0.76 – 1.24 | 0.813 |
| gibbous) |  |  |  |
| cond(LunarPhase8Waxing | 1.26 | 0.99 – 1.61 | 0.057 |
| crescent) |  |  |  |
| cond(LunarPhase8Waxing | 0.86 | 0.67 – 1.10 | 0.232 |
| gibbous) |  |  |  |
| N _DiverCode_ \| 45; N _SiteCode_ \| 17; Observations \| 29805 |  |  |  |

***Residual Plots***

*Whale Sharks:*

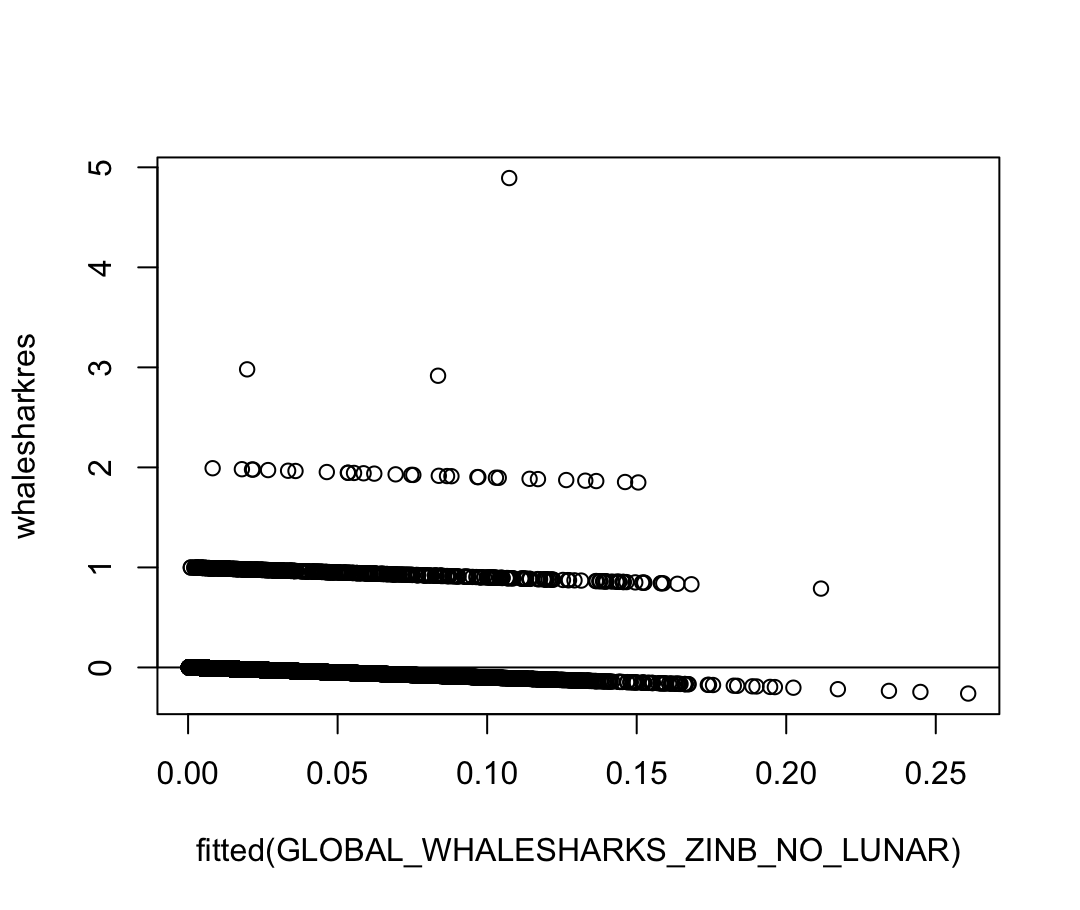

**Figure 1:** residual vs. fitted for whale shark top model

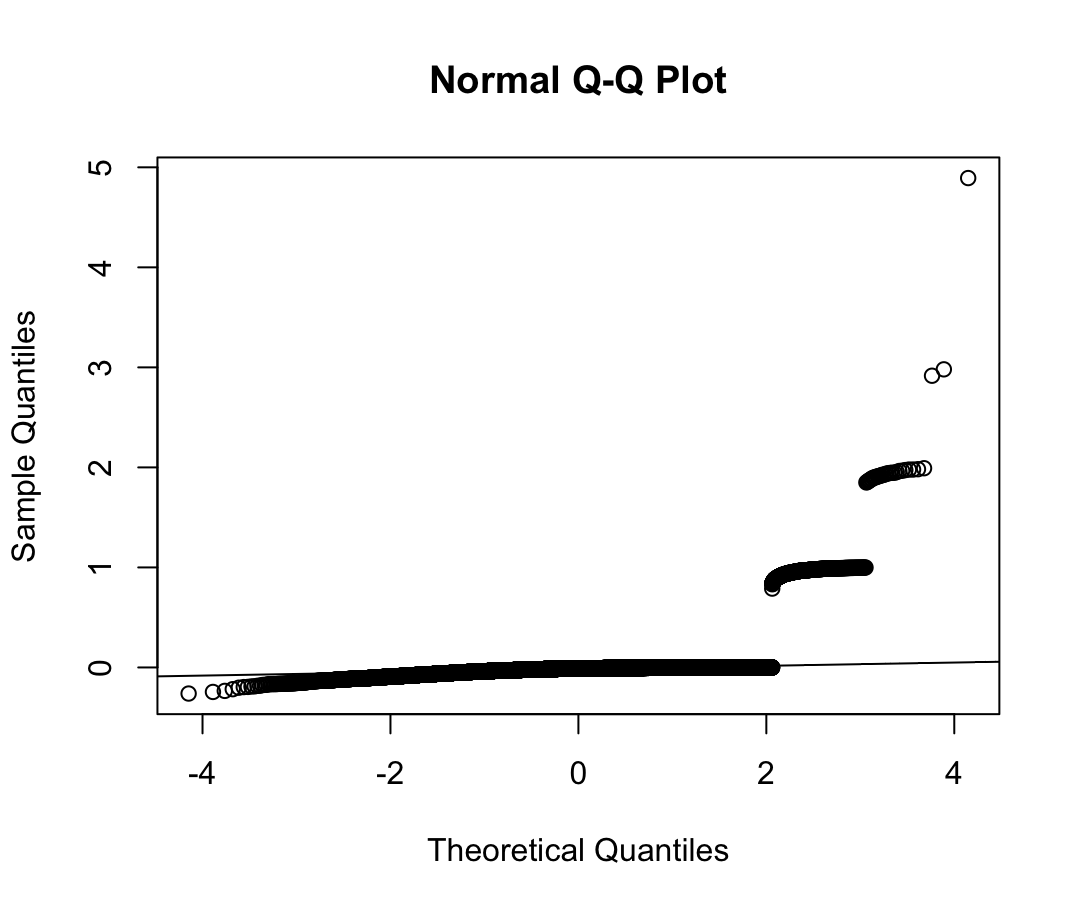

**Figure 2:** normal Q-Q Plot for whale shark top model

*Manta Rays:*

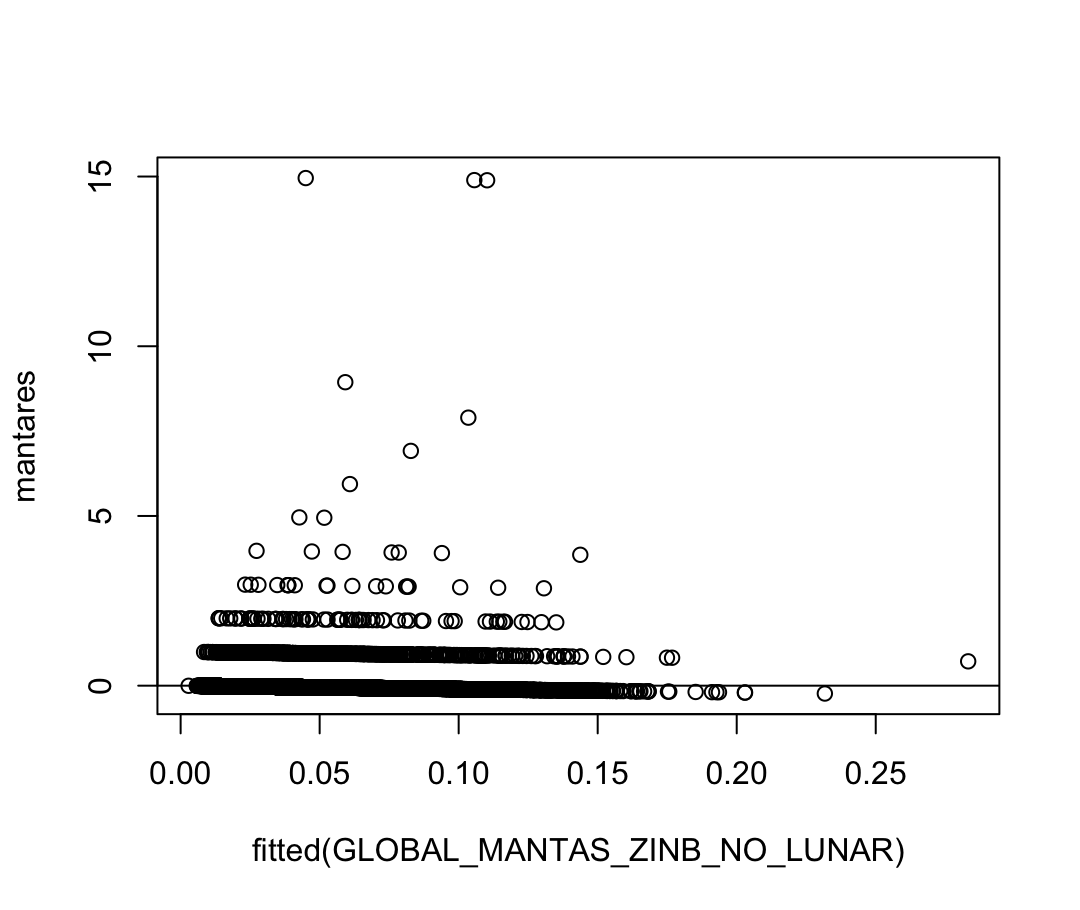

**Figure 3:** residual vs. fitted for manta top model

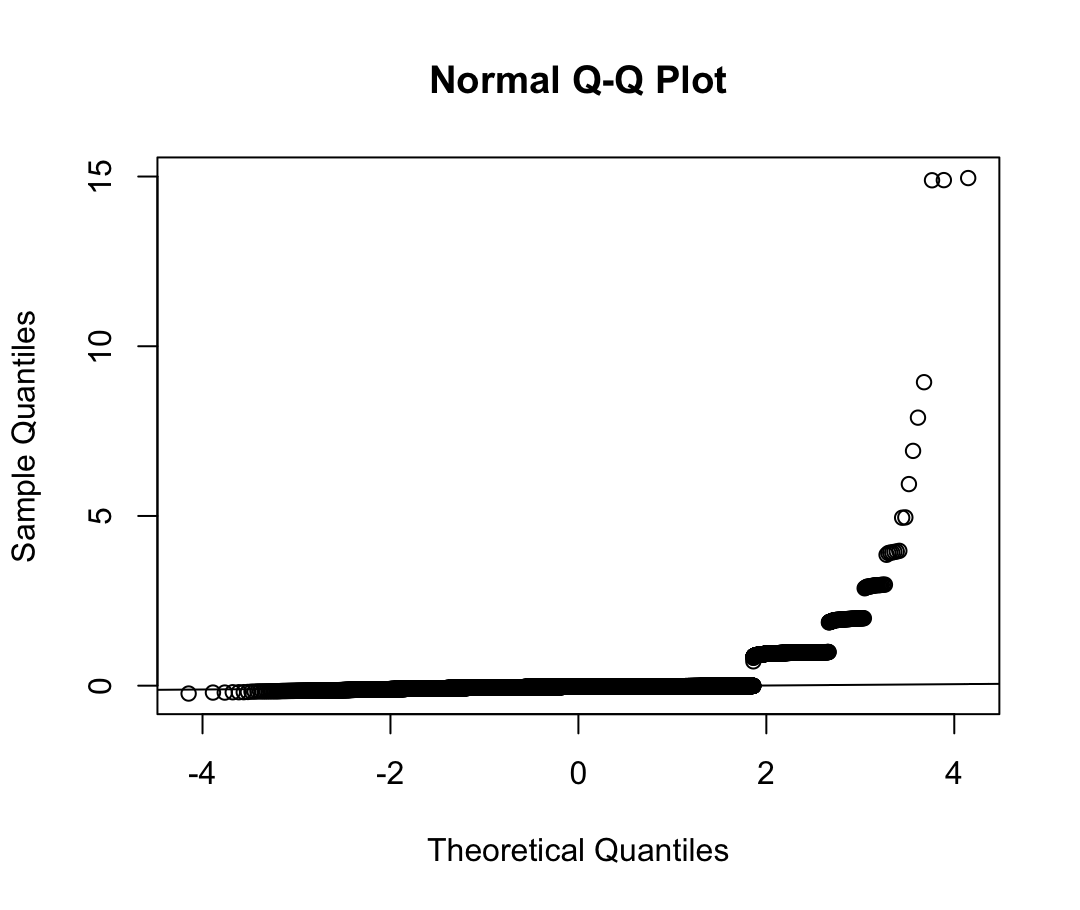

**Figure 4:** normal Q-Q Plot for manta top model

*Mobula Rays:*

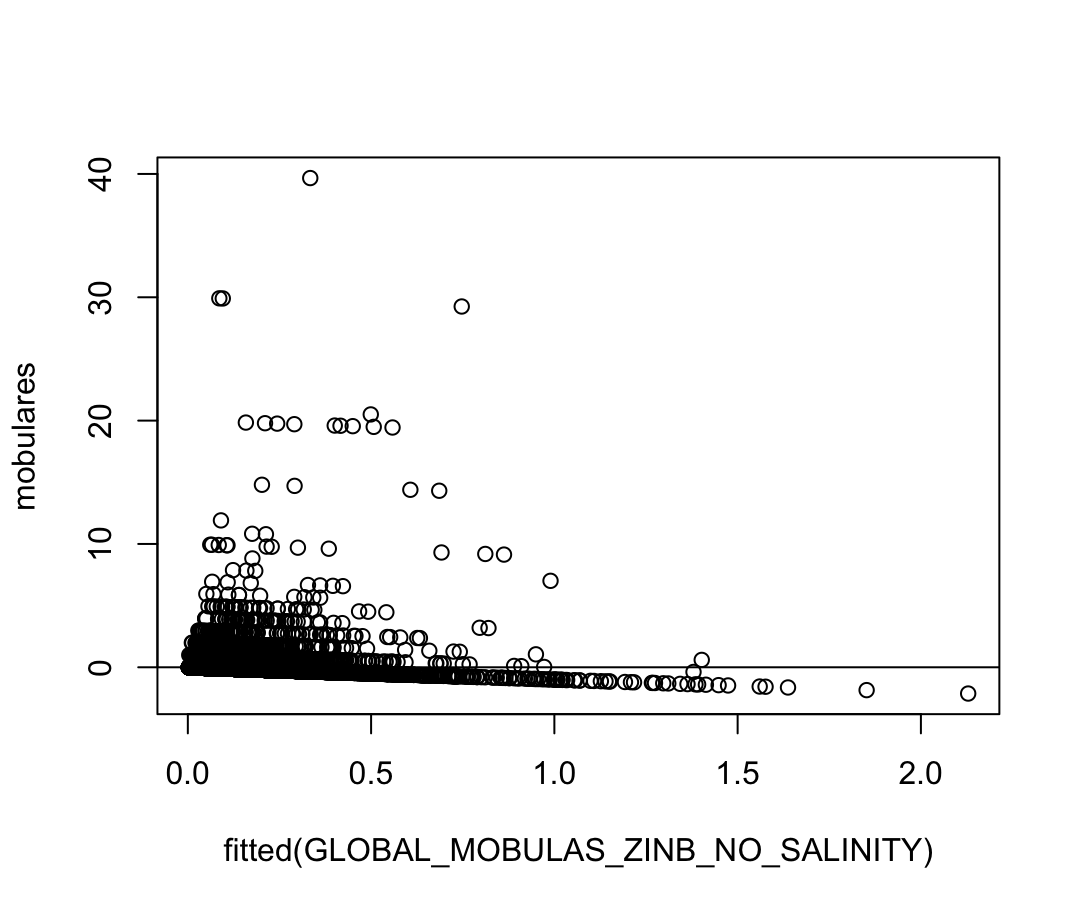

**Figure 5:** residual vs. fitted for mobula top model

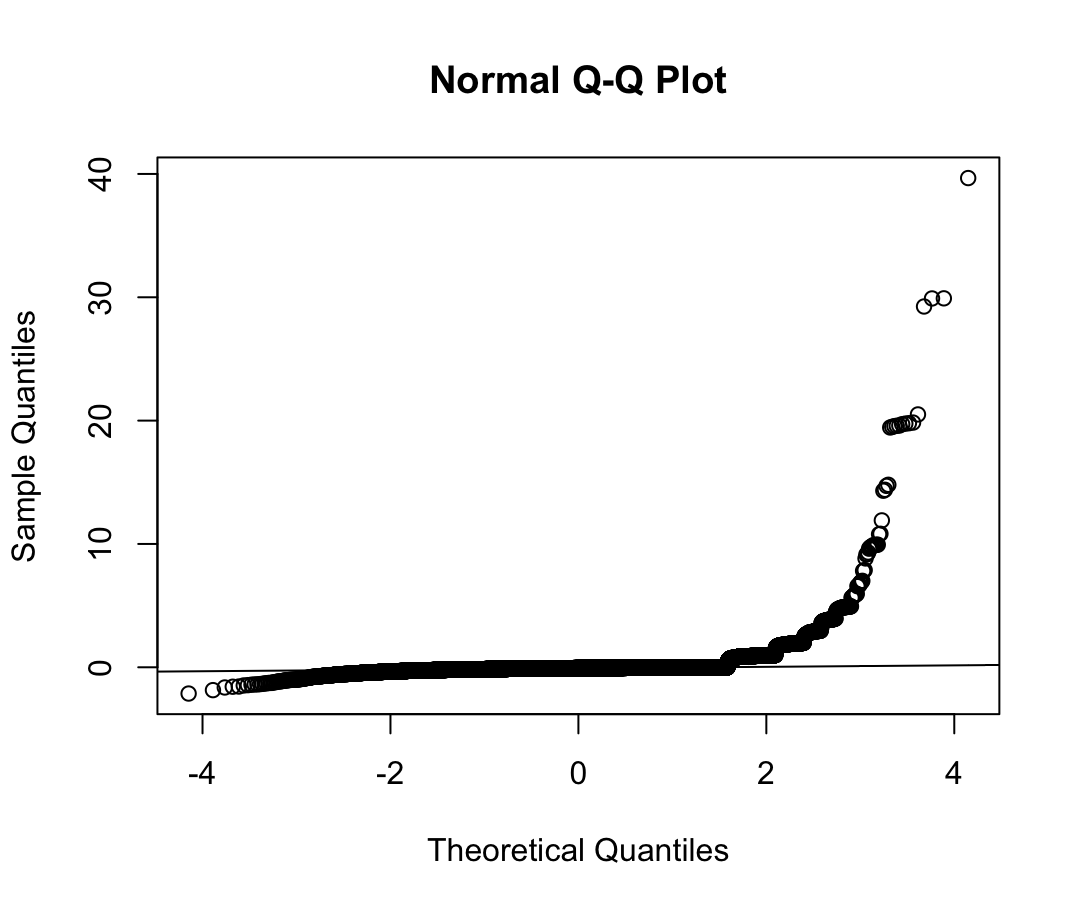

**Figure 6:** normal Q-Q Plot for mobula top model

***Supplemental Model Rankings and AIC***

**Table 4:** whale shark model rankings and AIC

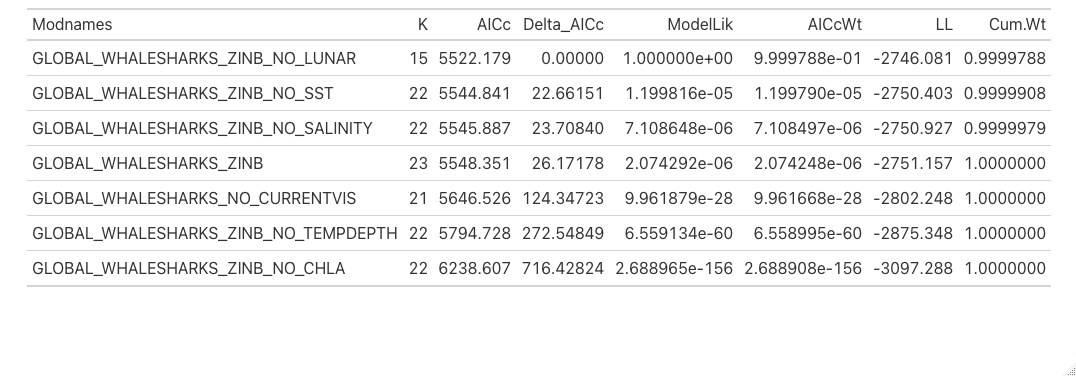

**Table 5:** manta model rankings and AIC

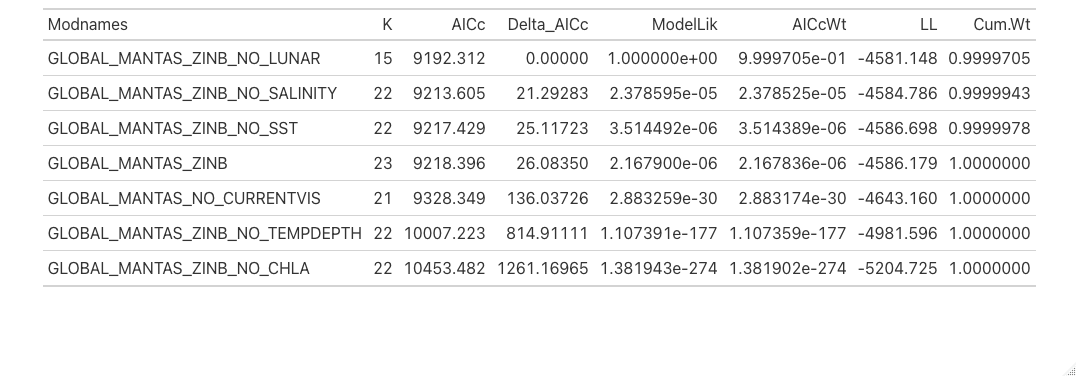

**Table 6:** mobula model rankings and AIC

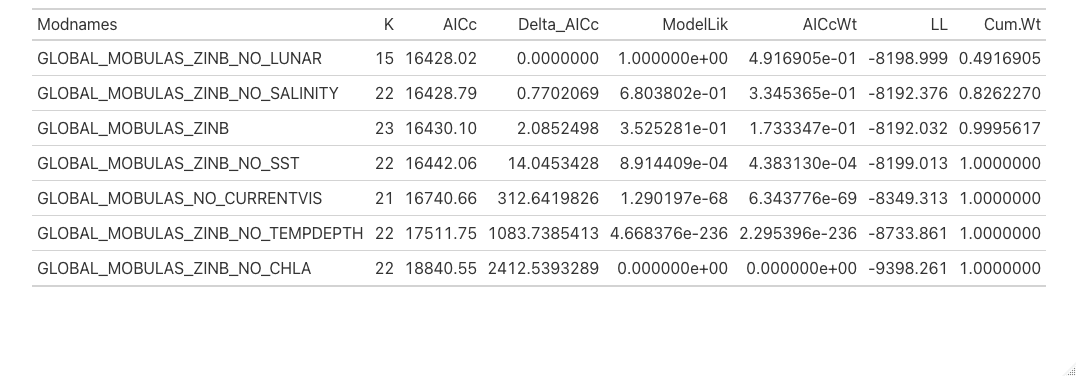

**Table 7:** Models used for AIC. Models were run for the presence and absence of the three species included in this study: mantas, mobulas, and whale sharks.

| **Model** | **Description** |
| --- | --- |
| Species Abundance ~ Ocean Nino Index + Temperature at Depth + Sea Surface Temperature + Salinity + Chlorophyll A + Lunar Distance + Lunar Phase + Current + Visibility + Year + sin () + cos() | Global Model |
| Species Abundance ~ Ocean Nino Index + Temperature at Depth + Sea Surface Temperature + Salinity + Chlorophyll A + Current + Visibility + Year+ sin() + cos() | Global Model without Lunar Factors |
| Species Abundance ~ Ocean Nino Index + Temperature at Depth + Sea Surface Temperature + Chlorophyll A + Lunar Distance + Lunar Phase + Current + Visibility + Year + sin() + cos() | Global Model without Salinity |
| Species Abundance ~ Ocean Nino Index + Temperature at Depth + Sea Surface Temperature + Salinity + Lunar Distance + Lunar Phase + Current + Visibility + Year + sin() + cos() | Global Model without Chlorophyll A |
| Species Abundance ~ Ocean Nino Index + Temperature at Depth + Salinity + Chlorophyll A + Lunar Distance + Lunar Phase + Current + Visibility + Year + sin() + cos() | Global Model without Sea Surface Temperature |
| Species Abundance ~ Ocean Nino Index + Sea Surface Temperature + Salinity + Chlorophyll A + Lunar Distance + Lunar Phase + Current + Visibility + Year+ sin() + cos() | Global Model without Temperature at Depth |
| Species Abundance ~ Ocean Nino Index + Temperature at Depth + Sea Surface Temperature + Salinity + Chlorophyll A + Lunar Distance + Lunar Phase + Year + sin() + cos() | Global Model without Current and Visibility |
